# Study on the Health Communication Behavior of Chinese Doctors on Network Media——Based on the Empirical Investigation of 119 Chinese Doctors

**DOI:** 10.1101/588863

**Authors:** Hu Fangyuan, Wang Pu

**Affiliations:** School of Journalism and Information Communication, Huazhong University of Science and Technology, Wuhan 430077, China

**Keywords:** Chinese Doctor, Network Media, Health Communication, Empirical Research

## Abstract

**Introduction:** In order to reduce the perceived risk of medical decision-making, Chinese patients tend to spend time and energy to understand relevant medical knowledge. After comparing online reviews and news reports of Chinese doctors, they carefully choose doctors. It is precisely because of the patient’s treatment habits that many doctors have spread health information through their own microblogs, WeChat public numbers, websites, videos and other means. Therefore, doctors’ speeches in cyberspace often play a dominant role in the development of health communication and guide the risk communication. This paper studies the influencing factors of Chinese doctors’ healthy communication behavior in online media. This has positive reference value for improving the health literacy of the whole people and the crisis management and guidance of public events.

**Methods:** In order to maximize the representativeness of the sample, the universality of the problems reflected in the survey data, and the adaptability of the purpose and method of the research questions, this paper adopts a multi-stage sampling method. In each region, a medical institution is selected according to a simple random sampling method in three types of hospitals: public hospitals, social hospitals, and primary medical institutions. A total of 10-11 medical institutions were selected in each region, and a total of 32 hospitals were selected from the three regions. According to the PPS sampling method, a corresponding number of working doctors are drawn in each hospital. Statistical analysis of the data was performed using SPSS 21.0 statistical software.

**Analysis:** The study found that doctors’ transmission of health-related content on online media is still at a lower level of involvement, and the frequency of transmission is significantly lower than the general level. There are also significant differences in the health transmission of doctors from different backgrounds on the Internet. In the context of control hospital background, continue to examine the impact of Media usage and occupational psychology on doctors’ dissemination of health-related content. The results show that the influence of Media usage on the health communication behavior of doctors on the Internet media, to a certain extent, verifies that the rise of online media plays a significant role in putting doctors into health science. This also shows that the media has fostered the doctor’s communication behavior. In addition, the higher the self-efficacy of doctors in the workplace, the more cautious they are about their words and deeds, the more they care about the impact of their speech on interpersonal communication.

## Introduction

On August 20, 2018, the Internet Information Center (CNNIC) released the 42nd “Statistical Report on the Development of China’s Internet Network” (http://www.cnnic.net.cn/). According to the data of the report, as of June 2018, the number of Internet users in China was 802 million, and the number of new netizens was 29.68 million, and the Internet penetration rate reached 57.7%. The number of mobile Internet users in China reached 788 million, and the number of new mobile Internet users was 35.09 million, an increase of 4.7% from the end of 2017. The proportion of Internet users using mobile phones has increased from 97.5% in 2017 to 98.3%, and the proportion of Internet users’ mobile Internet access continues to climb. This provides a reference for the public to seek medical information.

In general, in order to reduce the perceived risk of medical decision-making, patients tend to work hard to understand relevant medical knowledge, and then carefully select doctors after comparing online reviews and news reports^1–3^. As a result, this has also inspired many doctors to publish and widely disseminate health information through their own microblogs, WeChat public account, website, video and other means. The spread of health information by doctors on the Internet has a more complex impact on the development of health knowledge and the formation and evolution of risk paradoxes^4–7^.

According to “the 2017 China Health and Wellness Development Statistics Bulletin” (http://www.nhc.gov.cn/), by the end of 2017, the total number of health workers in China reached 11.749 million, including 3.39 million practicing doctors and practicing assistant doctors, and a total of 901,000 rural doctors. The number of practicing (assistant) doctors per thousand population reached 2.44, exceeding the international average. The development of the medical community has also led to an increase in the supply of medical resources. In 2017, the total number of medical practitioners in the national medical and health institutions was 8.18 billion, an increase of 3.2% over 2016. Among the doctors, the proportion of undergraduate and above has reached 39.76%, and the proportion of graduates or above is 3%. It can be seen that most doctors are subject to higher education, and their statements in cyberspace often play a dominant role in the development of health communication and guide the risk communication. Therefore, the study of doctors’ healthy communication behaviors on the online media has positive reference value for improving the health literacy of the whole people, the supervision of public opinion in the government, and the crisis management and guidance of public events^8–11^. In view of this, this study explores the following questions: 1. What is the current state of health communication among doctors on the Internet? 2. What is the difference between the health communication behavior of doctors at different levels of medical institutions on the online media. 3. What factors will affect the health communication behavior of doctors on the online media.

## Methods

All the samples in this survey were from the “Doctor Health Communication Behavior and Attitudes” questionnaire conducted in Wuhan hospitals from February 2018 to July 2018. The surveyed institutions included public hospitals, social private hospitals and grassroots hospitals. The target sample size of this survey was 150. After a six-month field survey, 119 valid questionnaires were collected, and the recovery rate was 79%. The specific survey methods are as follows:

### Survey area

The survey area is divided by the geographical distribution of the three towns of Hankou, Hanyang and Wuchang, each covering 1 to 2 municipal-level regional medical centers. In the areas of Optics Valley, Caidian, Huangpi and other areas, each area covers 1-2 comprehensive or specialist medical service institutions. Due to the wide coverage, the doctor’s survey sample of this study can be considered as a representative sample of doctors in central cities in central China.

### Hospitals surveyed

The total sample of this survey was derived from the list of medical institutions announced by the Wuhan Municipal Health and Family Planning Commission (http://www.whwsjs.gov.cn/) as of the end of December 2017. According to the data of Wuhan Medical Organization Planning Plan (2016-2020) formulated by Wuhan Municipal Health and Family Planning Commission, Wuhan Municipal Development and Reform Commission and Wuhan Municipal Finance Bureau, the city’s public hospitals accounted for 57.57%, social private hospitals accounted for 5.32% and grassroots hospitals accounted for 37.11%. In this survey, there are a total of 32 medical institutions in Wuhan, including public hospitals, social private hospitals, and grassroots hospitals.

### Sample extraction

The choice of sampling method for the study depends on the purpose of the study, the theoretical analytical framework used, the sample size of the survey, and the material conditions required for the survey. Based on this, in order to maximize the representativeness of the sample and the universality of the problems reflected in the survey data, and the suitability of the research objectives and methods, this study will adopt a multi-stage sampling method. The composition of the samples at each stage and the sampling degree are assigned as follows:

Sampling hospital. According to the data and level provided by the Wuhan Municipal Health and Family Planning Commission, as of the end of 2015, there were 5,341 medical institutions, including 285 hospitals. According to the nature of the organization, there are 113 public hospitals and 172 social private hospitals. There are 2,222 grassroots medical and health institutions in the city, including 133 community health service centers, 317 community health service stations, 69 township health centers, and 1703 village health clinics. There are 18 women and children health care institutions in the city; there are 2,816 other medical institutions such as outpatient departments and clinics in the city. In each area, according to three types of hospitals: public hospitals, social private hospitals, and grassroots hospitals, a medical institution is selected by simple random sampling. A total of 10-11 medical institutions were selected in each region, and a total of 32 hospitals were selected from the three regions.

### Sampling doctors

According to the pps (probability proportionate to size sampling) sampling method, a corresponding number of doctors are sampled in each hospital. When sampling doctors in 11 medical institutions in each region, because the total number of doctors in each region is inconsistent, the number of doctors sampled is calculated according to the proportion of doctors to be sampled in each region, that is, the number of doctors sampled / the total number of hospitals in the medical institution = Number of doctors to be investigated in the area / Total number of doctors in 11 medical institutions. The selection of specific sampling methods takes into account the number of medical treatments in this survey as many as 32. Some medical institutions have difficulty obtaining a full list of doctors to achieve random sampling, and also consider the feasibility of the survey. At the time of sampling the doctors, they were randomly selected at 9:00 in the morning when all the doctors in the department handed over the class.

### Statistical Analysis

Through the interviews with Chinese doctors, the close concerns of the influencing factors were collected. The interview results were summarized and summarized as the basis for the questionnaire design. This provides practical guidance for the research of the subject. Then, the survey was conducted by means of on-site questionnaires, and the data required for the research of this topic was collected. Finally, SPSS21.0 statistical analysis software is used to process the data. The analysis methods include descriptive statistics, single factor analysis and multiple linear regression analysis, and the statistical results are analyzed and discussed..

## Results

### The current status of doctors’ health communication on online media

The doctor’s health communication on the Internet media is measured by asking the doctor the frequency of posting his own science content on the online medium or to forward the frequency of health-related content. The results show that doctors who distribute a lot of their own science or forward health-related content on the network account for 1.7%, more accounted for 7.2%, generally accounted for 32.1%, relatively less accounted for 35.9%, almost never released or shared accounted for 23.1%. As shown in Figure 1, the frequency at which doctors post or share health-related content on the web media from “nearly released” to “frequently released” is assigned a value of 1-5. The statistics show that doctors post or forward health-related content on the network. The mean frequency was 2.34. The results of the one-sample T test showed that the frequency of healthy communication of doctors on the network was significantly lower than the general level (t=−48.795, p<0.05). It can be seen that the doctor’s healthy communication behavior on the network media is still in a state of low involvement.

**Figure 1.**
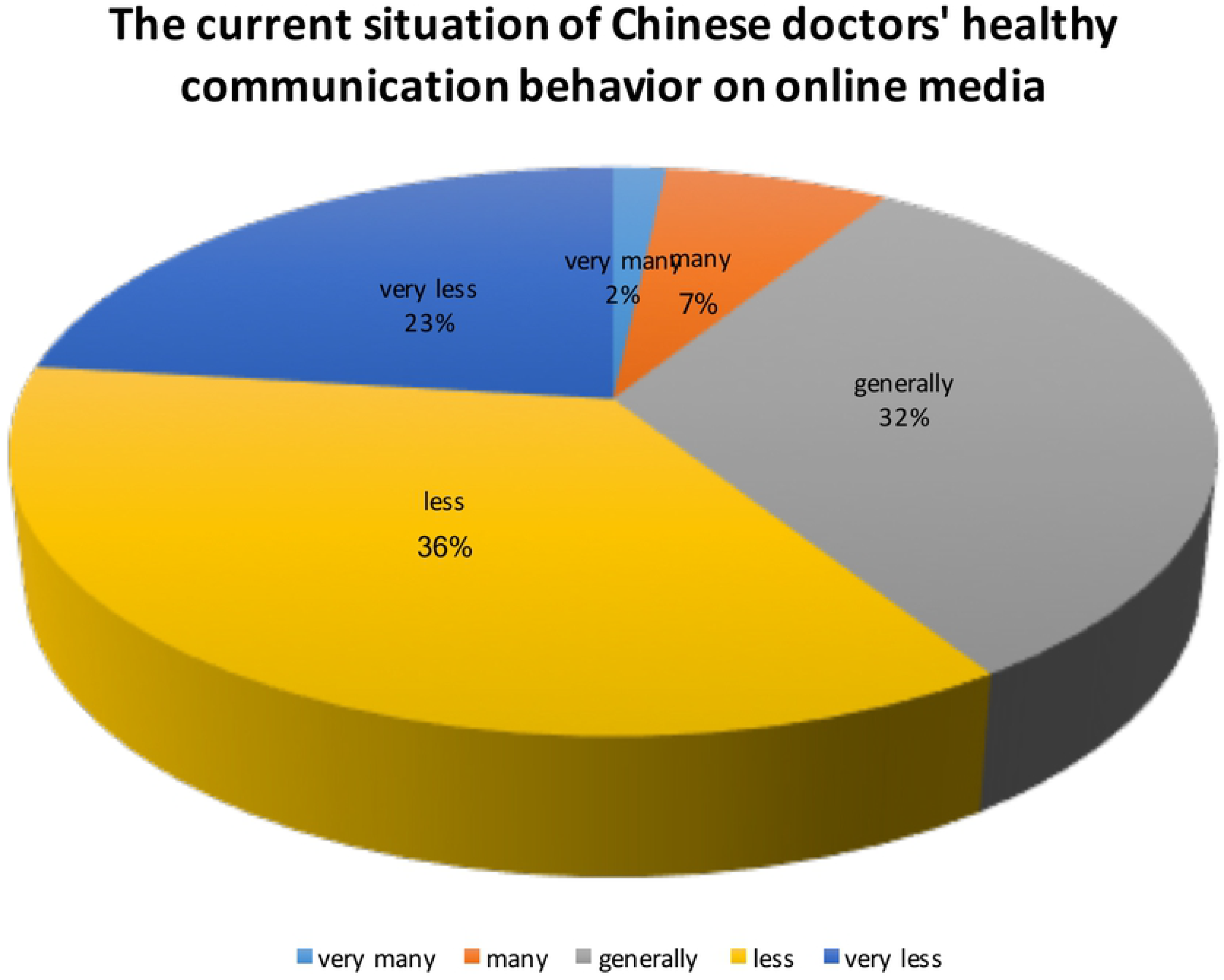

**Table 1.** Multi-level regression analysis of factors influencing the health communication behavior of Chinese doctors on online media

|  |  | Model<br>one | Model<br>two | Model<br>three |
| --- | --- | --- | --- | --- |
| Control variable | level one |  |  |  |
|  | gender | 0.131** | 0.124** | 0.125** |
| Background<br>variable | job title | 0.101*** | 0.099*** | 0.103*** |
|  | Department | 0.069** | 0.066* | 0.068* |
|  | Hospital type | 0.091*** | 0.088*** | 0.089*** |
| Media use | level two |  |  |  |
|  | Browse WeChat Duration |  | 0.145*** | 0.143*** |
|  | Browse Weibo Duration |  | 0.127** | 0.129** |
| Browse media | Browse Website duration |  | 0.108** | 0.106** |
|  | Browse network media<br>frequencies |  | 0.178*** | 0.179*** |
|  | Number of online friends |  | 0.189** | 0.187** |
| Media literacy | Media literacy |  | 0.101** | 0.102** |
| Professional attitude | level three |  |  |  |
|  | Career satisfaction |  |  | 0.096** |
|  | Self-efficacy |  |  | -0.055** |

Note: \*P< 0.05, \*\*P< 0.01, \*\*\*P<0.001

### Characteristics of health communication behaviors of doctors with different backgrounds on online media

A comparative analysis of the differences in behaviors of health-related content posted or forwarded by doctors of different backgrounds on a network medium. Regarding gender, the results of the independent sample t test showed that there was a significant difference in the healthy communication behavior between female doctors and male doctors on the Internet (t=−5.756, P<0.001). Specifically, the frequency of female doctors posting on the Internet or forwarding health-related content (M=2.32) was higher than that of male doctors (M=2.13). Regarding the professional title, the results of one-way analysis of variance were significantly different (F=9.571, P<0.001). Among them, the frequency of resident doctors publishing health-related content on the network media was the lowest (M=2.06), followed by the attending doctor (M=2.26) and the chief doctor (M=2.31), and the deputy chief doctor has the highest frequency of health communication on the Internet (M=3.36). Regarding the department, the results of the independent sample t-test showed that the doctor in the surgical department (M=2.29) was higher than the non-surgical department (M=2.11) (t=−3.526, P<0.001). This shows that doctors in the surgical department are more likely to conduct health communication on the Internet than doctors in the non-surgical department. Regarding the type of hospital, the results of one-way analysis of variance showed that doctors of different hospital types had significant differences in the frequency of dissemination of health-related content on the network (F=17.452, P<0.001). Specifically, the average frequency of health products distributed by public hospitals, social hospitals, and primary health care institutions on the Internet is 2.31, 2.15, and 1.93. It can be seen that hospitals in public hospitals are most likely to disseminate health-related content on the Internet.

### Influencing factors of doctors’ health communication behavior on online media

This section analyzes the influencing factors of doctors’ health communication in online media in three aspects: Media usage (including media trust, media literacy and media exposure) on the health communication of doctors on online media; the influence of social mentality on doctors’ health communication on online media; the impact of professional attitudes (including occupational satisfaction, professional trust, and professional efficacy) on the health-related content of doctors on the online media.

#### Media usage

Media usage usually refers to the frequency of use of the media by the audience, content preferences, and so on. The purpose of daily media use is to satisfy personal interests, increase knowledge, meet social, entertainment time, etc. ^12–14^. The purpose of daily Media usage is to satisfy personal interests, increase knowledge, meet social and entertainment time, and so on. The Media usaged in this study is divided into two parts: media exposure and media literacy.

Media exposure. First, analyze the correlation between the length of time that doctors browse online media such as WeChat, Weibo, and websites and the frequency of doctors’ health communication on online media. The results showed that the length of the length of time that doctors browse online media such as WeChat, Weibo, and websites was positively correlated with the frequency of doctors’ health communication on the network. The correlation coefficients were 0.195, 0.183, and 0.183, respectively, and the P values were less than 0.01. Secondly, analyze the correlation between doctors’ exposure to network media frequency, the number of online friends and the frequency of healthy communication behavior on the network media. The results show that the network media browsing frequency and the number of network friends are positively correlated with the frequency of posting or forwarding health-related content on the network medium. The correlation coefficients are 0.193 and 0.112, respectively, and the P values are less than 0.01. In other words, the higher the frequency of doctors’ contact with the network media and the greater the number of network friends, the higher the frequency with which doctors can post or forward health-related content on the network medium.

Media literacy. According to Rubin’s analysis, there are three main aspects of media literacy, namely, ability mode, knowledge mode and understanding mode^15^. As far as the ability model is concerned, it refers to the ability of citizens to acquire, analyze, evaluate and transmit various forms of information, focusing on the cognitive process of information; the knowledge model believes that media literacy is about how media contribute to society, with a focus on how information is transmitted. The understanding mode claims that media literacy is to understand that media information is subject to the coercive role of cultural, economic, political, and technological forces in the process of manufacturing, production, and communication, with a focus on the ability to judge and understand information. What is media literacy? It refers to people’s ability to choose, understand, question, evaluate, create, and think in response to various media information. The media literacy of this study was measured by asking doctors how they agree with the following options: I know how to publish my own health science or medical insights; I can identify the meaning and value of health information on the web; I know how to spread health knowledge on the web; I know how to use the medium to shape the image of a doctor. All of the above questions were evaluated and assigned using the Likert Level 5 scale from “1=nearly understood” to “5=very well understood”. The average score of these four questions was used to indicate the doctor’s media literacy (M=3.84, SD=0.612). The results of the correlation analysis showed that the doctor’s media literacy was positively correlated with the frequency of posting or forwarding health-related content on the online media (r=0.105, P<0.01). Therefore, the higher the doctor’s media literacy, the higher the frequency of posting or forwarding health-related content on the web media.

#### Social mentality

From the perspective of social psychology, social mentality is linked to specific social conditions or major social changes. In a certain period of time, it exists widely in the sum of emotions, social cognition, value orientation and behavioral intentions within social groups^16^. The social mentality of this study was measured by social sentiment. The doctor’s social sentiment is indicated by asking about the extent to which the doctor’s willingness to participate in society (from “1 = very unwilling” to “5 = very willing”). However, the results of the relevant analysis showed that the doctor’s social sentiment showed a negative correlation with the publication or forwarding of health-related content on the network, but it was not statistically significant (r=−0.012, P>0.05). But it also shows that doctors’ social sentiment does not affect their spread of health-related content on the online media.

#### Professional attitude

Occupation is not only a combination of the professional qualifications acquired by the individual and the acquired work experience. It is also a carrier of the integration of the individual and the society, a space of the essence of the interaction between the individual and the society, and the most essential of integration into society. All levels of the profession audiences’ demand and possession of social information in this media system appear in the form of a fusion and conflict. The audience and the media system are in an adaptive-balance-fusion interaction. Therefore, from the analysis of the current state of the doctor’s professional mentality, the relationship between the doctors’ health communication behavior on the network and the professional mentality media were analyzed and the degree of influence. The professional mentality of this study was explored through two dimensions: professional satisfaction and self-efficacy.

Professional satisfaction. The occupational satisfaction of doctors refers to the overall, emotional and emotional feelings and opinions of the occupations, working conditions and conditions that doctors are engaged in. It is also a kind of work attitude. The higher the doctor’s satisfaction with the profession, the more enthusiasm he or she will show to the profession, otherwise it will show complaints and indifference^17^. Therefore, doctors’ satisfaction with occupation may affect their initiative and enthusiasm for health communication. In this study, the Likert 5 scale was used to assign doctors’ professional satisfaction from “1=very dissatisfied” to “5=very satisfied”. The results of the correlation analysis showed that doctors’ satisfaction with occupation (r=0.051, P<0.01) was positively correlated with the frequency of healthy communication behavior on the network.

Self-efficacy. According to social cognition theory, self-efficacy has a significant predictive effect on human motivation, behavior, and attitude^18^. The professional efficacy of this study was measured by the doctor’s consent to the following statements: I am confident in my professional skills; the operation of the hospital is a complex system that I cannot understand; commenting on the Internet is one of the ways I express my views on hospital development.; I don’t think the hospital will take into account my opinions and ideas, from “1=very agree” to “5=very dissatisfied”. The average score of these four questions was used to indicate the doctor’s self-efficacy (M = 2.92, SD = 0.791). The higher the score, the higher the doctor’s self-efficacy. The results showed that the doctor’s self-efficacy was inversely related to the frequency of healthy communication on the network (r=−0.103, P<0.01). It also shows that the higher the self-efficacy of doctors in hospitals, the lower the frequency of dissemination of health-related content on online media.

### Regression analysis of the influencing factors of doctors’ health communication behavior on online media

The above correlation analysis is only a step-by-step test of factors that may affect the behavior of doctors on the Internet, but whether these factors will affect the health behavior of doctors on the Internet, as well as the intensity and direction of impact, further verification was performed using multiple regression analysis. Therefore, this study specifically explores the effects of doctor background, Media usage, and occupational psychology on doctors’ health communication behaviors on online media by establishing a multivariate hierarchical regression model.

In the regression analysis, the first layer inputs gender, title, department, and hospital type; the second layer inputs the media usage status, that is, media contact and media literacy; the third layer inputs professional attitude, that is, occupational satisfaction and self-efficacy. In the table, at the doctor’s background level, gender (β=0.131, P<0.01), professional grade (β=0.101, P<0.001), department type (β=0.069, P<0.01), hospital type (β=0.091), P < 0.001) has an impact on the behavior of doctors on the spread of health-related content on the Internet. Control the doctor’s background variables and continue to examine the impact of doctor Media usage.

Regarding the impact of Media usage, the online duration of the doctor on WeChat (β=0.145, P<0.001), Weibo (β=0.127, P<0.001) and website (β=0.108, P<0.001) has a positive impact on the doctor’s health communication behavior. This shows that the longer the doctors spend online on WeChat, Weibo and the website, the higher the frequency of healthy communication on the network. The health communication behavior of doctors on the online media was also affected by the online frequency (β=0.178, P<0.001) and the number of online friends (β=0.189, P<0.001). The higher the online frequency of doctors on the online media and the greater the number of online friends, the higher the frequency with which they post or forward health-related content on the network. Secondly, the media literacy of doctors also has a positive impact on the behavior of healthy communication on the network (β=0.101, P<0.001). The higher the media literacy of doctors, the higher the frequency of health communication on the network. Control the doctor’s background and Media usage variables to continue to examine the impact of the doctor’s social mentality.

Regarding the impact of occupational mentality, examining occupational satisfaction positively affected their behavior in healthy communication in the network (β=0.096, P<0.01). In other words, the higher the doctor’s satisfaction with the job, the higher the frequency of healthy communication on the network. Finally, the doctor’s self-efficacy negatively affected their healthy communication behavior on the network (β=−0.055, P<0.01). The higher the self-efficacy of doctors in the workplace, the less likely they are to spread health-related content on the online media.

## Discussion

Based on the questionnaire data of 119 doctors, this study explores the current status and influencing factors of doctors’ health communication on online media. The study found that doctors’ communication of health-related content on online media is still at a lower level of involvement, and the frequency of communication is significantly lower than the general level. There are also significant differences in the health communication of doctors from different backgrounds on the Internet. Specifically, female doctors may be more likely to communicate healthily on online media than male doctors; deputy director doctor-level doctors are more likely to disseminate health-related content; surgeons in surgical departments are more likely to use network media to spread health-related content than non-surgical doctors; public hospital doctors are more likely to disseminate health-related information on online media. In the context of control hospital background, this study continues to examine the impact of Media usage and occupational psychology on doctors’ dissemination of health-related content, with the following conclusions.

### Medium used

The length of time that doctors contact WeChat, Weibo and the website has a positive impact on the doctor’s health communication behavior. The longer the doctors contact WeChat, Weibo and the website, the higher the frequency of health communication on the network. The health communication behavior of doctors on the Internet is also affected by the frequency of contacts and the number of online friends. The higher the frequency of contact between doctors and the number of online friends, the higher the frequency with which they post or reprint health-related content on the network. This may be because when doctors browse online media such as Weibo, WeChat, and websites, there is an interaction between doctors and doctors, between doctors and patients, and it also mobilizes the enthusiasm of doctors for health communication. The intensity of the use of online media and the number of online friends have increased the spread of health-related content to a certain extent, providing greater possibilities for doctors to browse, acquire and disseminate health-related content.

The doctors’ media literacy also has a positive impact on the behavior of healthy communication on the online media. The higher the doctor’s media literacy, the higher the frequency of healthy communication on the network. Therefore, media literacy is also a key media factor that affects doctors’ dissemination of health content.

In short, the influence of Media usage on the health communication behavior of doctors on the Internet media to some extent verifies the shaping effect of the rise of online media on doctors’ investment in universal health science, and emphasizes the cultivating effect of media on doctors’ expression and dissemination.

### Professional attitude

The occupational satisfaction of doctors is positively affecting their behavior in the online media. The higher the doctor’s satisfaction with the profession, the higher the frequency of healthy communication on the network. The self-efficacy of doctors is negtively affecting their healthy communication behavior on online media. This also shows that the higher the self-efficacy of doctors in the workplace, the more cautious about their words and deeds, the more concerned about the impact of their own speech on the acquaintance circle.

However, this study also has some shortcomings. When examining social mentality and professional mentality, there is a lack of multiple dimensions. Future research should examine doctors’ social mentality and professional mentality from multiple dimensions to better explain the influence of social mentality and professional mentality on their communication behavior. However, this study also has certain theoretical and practical significance: it theoretically proves the influence of doctor background and Media usage on doctors’ dissemination of health-related content on online media, and also provides empirical support for the popularization of national health knowledge and the guidance of risk public opinion.

